# Anticipated Relevance Prepares Visual Processing for Efficient Memory-Guided Selection

**DOI:** 10.1101/2025.08.20.671232

**Authors:** Lasse Dietz, Christoph Strauch, Kabir Arora, Stefan Van der Stigchel, Samson Chota, Surya Gayet

## Abstract

Finding an object typically involves using working memory to select relevant visual information *at the right time*. Using EEG, we investigated how predictable changes in stimulus relevance influence top-down visual selection. Participants memorized an oriented grating, followed by a cue indicating which of two sequentially presented probes was relevant for a memory-match/-mismatch judgment. Consistent with earlier work, relevant probes evoked stronger univariate responses than irrelevant probes. Furthermore, responses to (memory-matching and memory-mismatching) probes were more distinct when they were relevant compared to irrelevant. Crucially, using rapid invisible frequency tagging (RIFT), we found that visual responses to the (empty) probe location were enhanced even before the presentation of relevant (compared to irrelevant) probes. Predictable changes in stimulus relevance, therefore, induce both pro-active and re-active changes in visual processing. We conclude that anticipating the (ir)relevance of upcoming visual events enables the visual system to prepare for efficient memory-guided visual selection.

## Introduction

Picture yourself driving on the highway and seeing a blue sign announcing your vacation destination, the city of Utrecht, in 5km. Because you roughly know how much time it takes to reach this exit, the immediately upcoming exit signs are irrelevant to you. Instead, you start to look for the blue highway exit sign to Utrecht after a short wait, find your exit, and enjoy your holiday.

In this example, selecting the relevant road sign is easy as you already know what to look for. Visual working memory (VWM) facilitates this selection process. VWM prioritizes relevant visual input by enhancing the processing of VWM-matching visual events (here: a blue sign) (Olivers et al., 2011; Soto et al., 2008). More specifically, VWM content modulates neural responses (Gayet et al., 2017), conscious access (Gayet et al., 2013), attentional capture (Downing, 2000), and gaze biases (Olivers et al., 2006; Olmos-Solis et al., 2017; Silvis & Van Der Stigchel, 2014) to VWM-matching compared to mismatching visual input. By doing so, VWM allows observers to guide visual selection toward currently relevant visual events (Desimone & Duncan, 1995; Duncan & Humphreys, 1989; Eimer, 2014; Kastner & Ungerleider, 2001; Wolfe, 1994; Wolfe & Horowitz, 2004).

But what will be relevant later may not yet be relevant now. In our highway example, the to-be-selected visual event (i.e., the exit sign) only became relevant later. Anticipating *when* relevant visual events will appear would allow observers to distribute cognitive resources across time efficiently. Indeed, temporal expectations can aid in perceptual discrimination and/or detection; successfully anticipating visual events is associated with altered neural responses and improved behavioral performance (Bertelson & Boons, 1960; Coull & Nobre, 1998; Cravo et al., 2013, 2017; Denison et al., 2024; Klemmer, 1956; Miniussi et al., 1999; Nobre & Van Ede, 2018; Rohenkohl et al., 2012; Vangkilde et al., 2012). Similarly, memory-guided (i.e., VWM-based) visual selection may benefit from anticipating the appearance of relevant visual events. As our highway example illustrates, memory-guided visual selection ("what are we looking for?") in daily life is strongly dependent on temporal expectations ("when are we looking for it?"). Thus, temporal expectations may profoundly influence working memory-guided visual selection as well.

Initial research studying anticipation in the context of memory-guided behavior shows that participants are more accurate (and faster) in change-detection tasks when the probe appears after the anticipated interval (Heuer & Rolfs, 2022; Jin et al., 2020). These findings show that temporal expectations improve performance in memory recognition tasks. Such improvements may be the result of either (A) pro-actively enhancing the representation of the memory content, or (B) improved sensory processing of the probe item (or both). Following the discovery that active VWM content can be decoded from fMRI data (Harrison & Tong, 2009; Serences et al., 2009), numerous studies have shown that observers can indeed alter the representational strength of VWM content over time, depending on when the recall task is expected to take place (Christophel et al., 2018; LaRocque et al., 2017; Lewis-Peacock et al., 2012; Rose et al., 2016; Sahan et al., 2020; van Loon et al., 2018; Yu et al., 2020). Converging evidence was found using a variety of methods, including pupil size measurements (Zokaei et al., 2019), directional microsaccade biases (Olmos-Solis et al., 2017), and alpha-band suppression in EEG recordings (Van Ede et al., 2017). In the auditory domain, temporal expectations were found to protect VWM content against temporal decay (Wilsch et al., 2018). Thus, it is well established that observers can alter the representational strength of VWM content in anticipation of task requirements. It remains unknown, however, whether visual processing is altered in anticipation of forthcoming sensory events that are relevant to memory-guided behavior. The reason for this is two-fold. Firstly, it is inherently challenging to measure visual processing in anticipation of (i.e., before the presentation of) visual events, because that period of interest (by definition) has no visual event to measure a response to. Secondly, existing neurophysiological markers that are often used to index anticipatory effects are not specific to visual processing (e.g., contingent negative variation, or CNV (Walter et al., 1964)) and/or their role in visual processing is not well understood (e.g., endogenous neural oscillations, such as alpha-band activity) and therefore may be contaminated by VWM-related processes, task difficulty, or motor preparation.

Here, we leveraged a novel EEG-based method that allows for continuous tracking of early visual processing in the absence of any perceptible stimulation. In doing so, we directly tested whether visual processing is enhanced in anticipation of visual events that are relevant (versus irrelevant) to memory-guided selection.

Participants memorized a single oriented grating on each trial, followed by a cue. The cue indicated which of two subsequent gratings had to be compared to the initial memory item (relevant probe) and which could be ignored (irrelevant probe; see Figure 1 for the paradigm). This design allowed us to investigate (1) how the anticipation of relevant versus irrelevant visual events affects visual processing pro-actively (before stimulus presentation), and how (2) the memory content influences the response to relevant versus irrelevant visual events re-actively (upon stimulus presentation). Importantly, relevant and irrelevant probes were physically identical across trials and only differed in terms of task-relevance, since stimulus changes (cue, orientation, etc) were counterbalanced within participants. The present design deviates from classical research on attentional selection processes, which typically involves multiple spatial locations. Such task designs prompt participants to prioritize specific spatial locations over others. In contrast, this task design only involves a single spatial location, with relevance varying over time. This approach uniquely allows us to measure changes in visual processing per se, separating them from local changes in visual processing caused by the reallocation of spatial attention.

**Figure 1:**
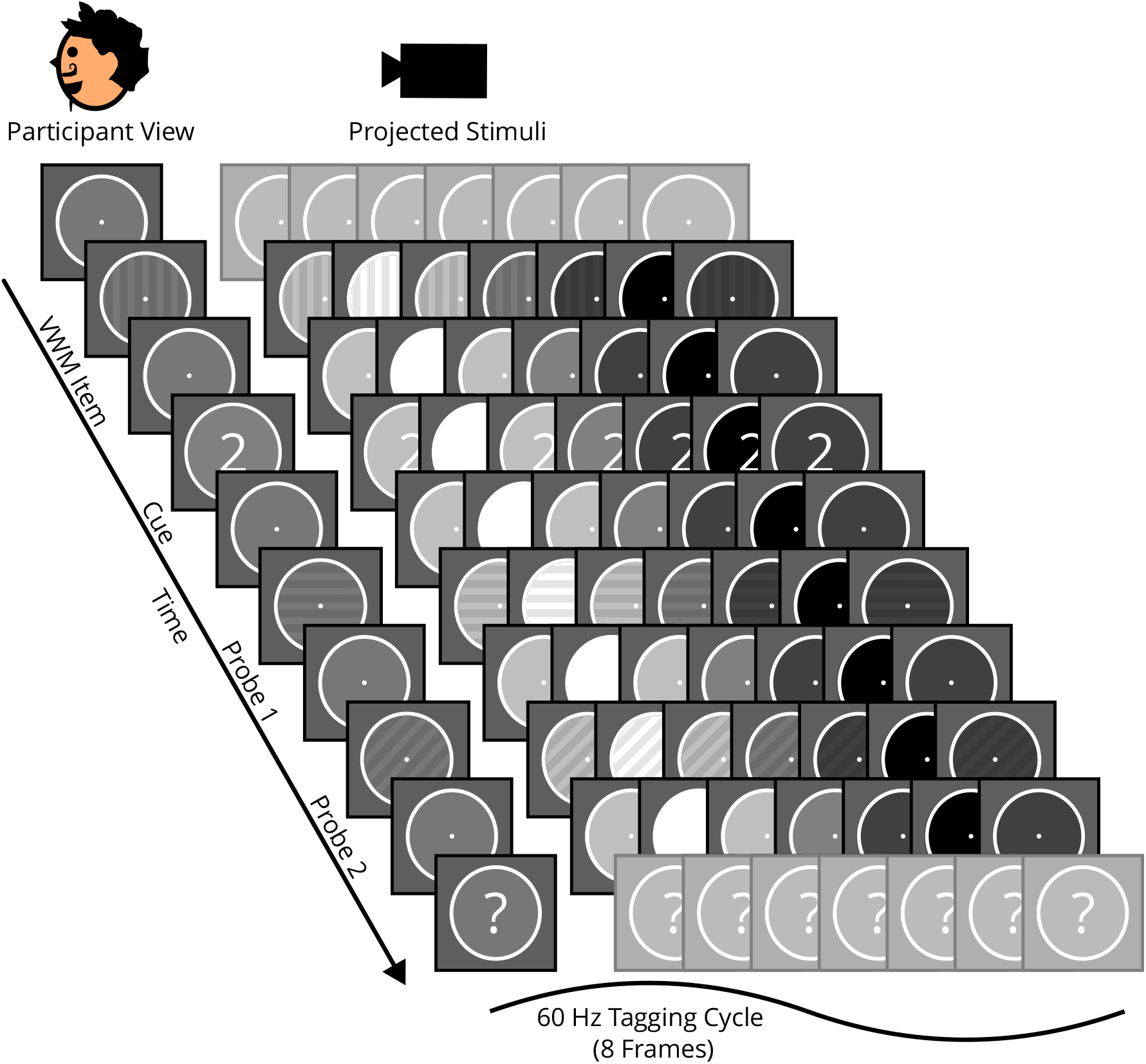
The left side displays the visual events as perceived by the participants. The luminance changes of a single RIFT cycle (8 frames including the left side) for each of these events are depicted horizontally. Luminance changed periodically at 60Hz, rendering the luminance changes imperceptible. The first fixation period and the response period are not tagged (continuous gray background). This is an example of a cue "2" trial (participants have to compare the memory item to the second probe) requiring a "different" response (the orientation of the second probe differs from that of the memory item). Untagged periods (fixation before the VWM Item and the response period) are grayed out.

To measure changes in early visual processing evoked by the mere anticipation of relevant and irrelevant visual events, EEG-based rapid invisible frequency tagging (RIFT) was employed (Arora et al., 2024; Hustá et al., 2025). RIFT can continuously track changes in early visual processing, even in the absence of any perceptible stimulus. This is achieved by ’tagging’ a location of interest on the screen (here: the stimulus area). Tagging refers to periodic luminance modulation of the screen at 60Hz (imperceptible to participants). This luminance modulation induces a corresponding oscillatory response in the EEG signal. The strength of this oscillatory response in the EEG (measured as RIFT coherence) varies with how much cognitive resources (attention) are allocated to visual processing of the tagged location (Arora, Gayet, Kenemans, et al., 2025; Çelik et al., 2025; Duecker et al., 2024; Ferrante et al., 2023; Wang, Arora, et al., 2025; Zhigalov et al., 2019). Crucial to the present purpose, RIFT produces an experimenter-induced marker of early visual processing, which enables us to isolate pro-active effects on visual processing that are not triggered (or modulated) by the conscious perception of a stimulus. In contrast, typical approaches using visible probes inevitably confound pro-active effects with re-active effects, which may not have emerged without said probes. Other approaches using endogenous markers of visual processing (e.g., alpha oscillations) cannot be unequivocally attributed to early visual processing, because they are not experimentally induced (’tagged’) and may therefore reflect a variety of (early and late, perceptual and post-perceptual) processes. To summarize, RIFT is a non-invasive, location-specific, experimenter-induced oscillatory signal with high temporal resolution that tracks early visual processing at the tagged location.

We predicted that if observers indeed (pro-actively) enhance sensory processing in anticipation of relevant visual events during memory-guided selection, the RIFT response should increase before presentation of a relevant probe compared to an irrelevant probe.

## Methods

### Experimental Model and Study Participant Details

Twenty-five participants with normal or corrected-to-normal vision were recruited at Utrecht University. Each participant attended a 2-hour session and was compensated with either participation credits or €24. Informed consent was obtained, and participants provided their age and gender. The study was conducted in accordance with the protocol approved by the Ethics Committee of the Faculty of Social and Behavioural Sciences at Utrecht University with reference number 863732 – HOME-OSTASIS (approved on 18/03/2020). Out of the initial 25 participants, two exhibited significantly poorer behavioral performance on either Cue 1 or Cue 2 trials (Cue 1: 48.61% versus Cue 2: 96.53% and Cue 1: 94.44% versus Cue 2: 62.50%). As these participants did not seem to adhere to the task instructions (they likely ignored the cue information), they were excluded from further analysis. Additionally, one participant was excluded due to poor EEG data quality. Consequently, a total of 22 participants were included in the eventual analyses.

### Method Details

#### Setup & Apparatus

Stimuli were created using Psychtoolbox for MATLAB (Brainard & Vision, 1997; Pelli, 1997) and presented on a projection screen using a ProPixx projector (VPixx Technologies Inc., QC Canada; resolution = 960×540px; refresh rate 480Hz). The projected screen size was 48×27.5cm or 35.05×20.51 degree visual angle (dva) in a rear-projection format. Interfacing with the projector (e.g., turning on/off 480 Hz mode) was achieved using the custom Matlab software provided by the company, which was integrated into the experimental code. The ProPixx projector was gamma-calibrated to a linear input-output function. Luminance as measured in a dark room using an external photometer (Gossen Mavo-Monitor USB) yielded 0.95cd/m2 for black (RGB [0, 0, 0]), 649cd/m2 for gray (RGB [127 127 127]), and 1330cd/m2 for white (RGB [255 255 255]) displays. Participants were positioned 76 cm from the screen, using a chin and headrest for alignment.

#### Task & Procedure

Participants performed a modified delayed match-to-sample task (see Figure 1, "Participant View"). Each trial began by pressing the space bar, which initiated a 1500ms baseline of untagged fixation, followed by a first oriented grating (hereafter: the memory item), which was presented for 200ms. Participants were instructed to memorize the orientation of this memory item for a subsequent match/mismatch judgment (i.e., they reported whether the cued probe had the same or a different orientation as the memory item). After a second fixation period (1000ms), a cue was presented for 200ms ("1" or "2") informing participants which of two subsequent oriented gratings (the probes; each presented for 200ms, with a 1500ms inter-stimulus interval) was relevant for the memory match/mismatch judgment. After a 500ms fixation period following the offset of the second probe, a question mark was presented, indicating the forced-choice response window (1000ms). Participants indicated whether the orientation of the cued probe (i.e., first or second) matched the orientation of the memory item by pressing either the left or right control key with the corresponding hand. Each trial ended with a blue circle, signaling a self-paced break. Participants were asked to refrain from using strategies to circumvent the memory component of the task. As an example, we specifically prohibited using one’s fingers as a memory aid for stimulus orientation. Participants were instructed that the irrelevant (i.e., uncued) probe could be ignored.

#### Stimuli

The stimuli were oriented square-wave gratings (diameter: 6 dva). Twelve orientations, ranging from 10 to 190°to avoid fully vertical and horizontal orientation in 15°steps, were randomly selected. The gratings consisted of alternating shades of gray (127.5 RGB & 95% of 127.5 RGB). The phase value was a random integer between -180 and 180, excluding 0, to prevent fixation on the stripes. The background color was a darker gray (100 RGB) to reduce eyestrain. Fixation dot, response indicator, and stimulus area frame were white (200 RGB). The gratings had soft borders (2D-Gaussian transparency mask) to prevent edge-based orientation memorization or recall strategies. 60Hz RIFT was achieved by multiplying the stimulus area RGB values with eight discrete steps of a sinusoidal curve from 0 to 1. A complete tagging cycle takes ∼16.667ms, which corresponds to a frequency of 60 Hz (Figure 1 - Projector view). This caused the lighter parts of the grating stimuli and the framed stimulus area to vary fully from 0 to 255 RGB, while the darker parts of the grating stimuli varied from 0 to 95% of 255 RGB. RIFT started at the onset of the memory item and ended at the onset of the response screen. Consequently, each trial had a period of 5300ms of uninterrupted 60Hz tagging.

#### Design

Participants completed 12 blocks of 24 trials each, lasting approximately one hour in total, excluding one practice block. The additional practice block consisted of 12 trials, with feedback provided for the first six trials. Per two blocks, trials were balanced for relevant/irrelevant (Cue 1/2) probe, the 12 orientation conditions (10° to 190°), and presented in random sequence. Probes were rotated in one-third of the trials (equally 45° to the right, left, or by 90°). Orientations of all gratings were independent of each other. Each memory item orientation appeared four times per two blocks. Trials within each set of two blocks were presented in a different random order per participant. At the start of a block, participants were informed about the current block number and afterwards about their accuracy in the past block for 3000ms.

#### Eyetracking

The eyes were binocularly recorded at 500Hz using an Eyelink 1000 eye tracker (SR Research, Ontario, Canada), with nine-point calibration and validation performed at the start and after each block.

#### Invalid Trials

If participants failed to give a response within the forced-choice window, the response was considered invalid. To ensure adherence to central gaze, trials were aborted and considered invalid if the gaze was outside the stimulus area for more than 208ms (25 frames at 120Hz), thus allowing a few short blinks. Blinks were monitored, using the live eyetracking data to detect NaN values. Participants were reminded before the next trial on screen to avoid blinking during trials. If invalid, trials were repeated up to two times at the end of blocks in a random sequence.

### Quantification and Statistical Analysis

#### EEG Data Collection & Preprocessing

EEG data were collected using a 64-channel ActiveTwo BioSemi system at 2048Hz. Preprocessing was done using the Fieldtrip toolbox for MATLAB (Oostenveld et al., 2011). Data was re-referenced to the average of all electrodes, line noise removed (50, 100, 150Hz) using a discrete Fourier transform, high-pass filtered (0.01Hz), and split into trials of 6800ms (1500ms before VWM onset & 5300ms until response screen start). Independent component analysis was used to remove oculomotor artifacts. Trials containing other artifacts were identified using visual inspection and removed (average trials excluded per participant = 40.41, *SD* = 26.62).

#### RIFT - Data Analysis

Magnitude Squared Coherence (as described in Pan et al., 2022) was used to quantify the strength of the EEG responses at our tagging frequency (60Hz). To this end, the EEG trials were band-pass filtered (±1.9Hz) at 60 Hz using a two-pass Butterworth filter (4th order, Hamming taper). The filtered data were then Hilbert transformed to obtain the analytic signal, which was used to calculate time-varying coherence:

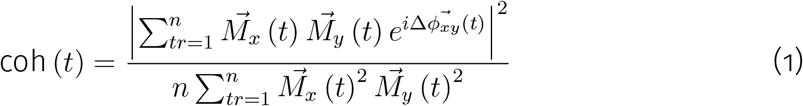

With n being the number of trials. For each time sample t in trial tr, the time-varying magnitude of an EEG signal is represented by 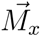 and by 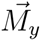 for a synthetic sinusoidal reference signal. The phase difference as a function of time is defined by 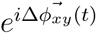. Coherence, therefore, reflects the degree to which the 60 Hz phase (weighted by magnitude) is consistently aligned across trials for each channel. The 60Hz coherence traces per condition (Cue 1, Cue 2) were averaged across each participant’s top five RIFT coherence channels. The top five channels were identified per participant by averaging RIFT coherence across all conditions (Cue 1 and 2) and time points throughout a trial. Previous literature indicates that the number of top channels included does not impact the results (Arora, Gayet, Kenemans, et al., 2025). Subsequently, per participant, the difference between the Cue 1 RIFT trace and the Cue 2 RIFT trace was calculated to reveal changes in visual processing for the two probe intervals.

#### Univariate Analysis

The preprocessing procedure was the same for all univariate analyses. To preprocess data for univariate analysis, data were downsampled to 128 Hz and frontal channels were removed (FT7, T7, FT8, T8, AF7, AF3, AFZ, AF4, AF8, F7, F8, FP1, FP2, FPz). The data were baseline corrected by subtracting the average of the baseline window (-600 to 100ms, before probe onset) from each trial per channel. The EEG data were segmented around probe onset from -650 to 1200ms (Probe 1) and from -650 to 700ms (Probe 2; I.e., until response onset).

To confirm that participants engaged with the temporal structure of the task, we compared univariate EEG responses to relevant versus irrelevant and memory-matching versus memory-mismatching visual events. Previous research suggests that responses to relevant (i.e., prioritized) compared to irrelevant visual events should be stronger in posterior channels (parietal/occipital) (Correa et al., 2006; Hillyard et al., 1998; Mangun & Hillyard, 1990; Miniussi et al., 1999). To test for the effect of relevant versus irrelevant visual events on the univariate EEG signal, we averaged EEG data across three evenly spaced time windows (200ms) starting at probe onset. We chose this data driven approach because we had no prior expectation for the topographical location of the effect of relevance. Within each window and for every channel, the response difference between relevant and irrelevant visual events was assessed using a two-sided t-test (p-values adjusted for multiple comparisons by controlling the false discovery rate (FDR; Benjamini & Hochberg, 1995; Benjamini & Yekutieli, 2001, 2005)).

Furthermore, if observers made effective use of the temporal expectations during memory-guided visual selection, the neural response difference between memory-matching and memory-mismatching visual events should be stronger when events are relevant compared to irrelevant. We leveraged the same approach as described above; Here, however, we compared the effect of memory-matching versus memory-mismatching visual events when they are relevant and when irrelevant (see Figure 3A). Because this approach is agnostic about the spatial location of the effect (but lacks temporal resolution), it allowed us to determine the shared set of channels between Probe 1 and Probe 2 capturing the difference between memory-matching and memory-mismatching responses for further analysis. For this channel selection, we only considered the memory-match versus memory-mismatch effect when the probe was relevant, because then the effect should be the strongest (i.e., the effect should be weaker/absent when probes are irrelevant). Based on this shared set of channels (marked with black asterisks in Figure 3A, middle row), we averaged across participants split on the conditions of interest (relevant & irrelevant, memory-match & memory-mismatch; see Figure 3B). The increased temporal resolution of the ERPs allowed us to employ tests with increased sensitivity (i.e., permutation cluster tests; see Significance testing) to test the four key comparisons:

1. Neural responses to relevant versus irrelevant visual events
2. Neural responses to relevant memory-matching versus memory-mismatching visual events
3. Neural responses to irrelevant memory-matching versus memory-mismatching visual events
4. Neural responses to relevant versus irrelevant memory-matching versus memory-mismatching visual events

#### Significance Testing

Significance for all time-re analyses was determined using non-parametric cluster-based permutation tests (see Maris & Oostenveld, 2007). The testing procedure consisted of three steps: 1) For each time point of the to-be-tested trace, a t-test identified samples differing significantly from zero (one-sided, *p*< 0.1). For clusters of at least 5 consecutively significant samples, the sum of t-values was computed, resulting in a cluster-level-t-mass. 2) To produce a null-distribution of t-masses (reflecting the expected distribution under the null hypothesis), all traces were randomly sign-reversed (preserving autocorrelations between time points in the null distribution) 5000 times, resulting in 5000 new datasets containing a randomized trace per participant. The same analysis procedure used for the observed data (Step 1) was applied to each of these 5000 datasets; except, only the largest cluster identified in each of the datasets was included in the null distribution. 3) t-mass values of the observed clusters higher than 95% (i.e., α = 0.05) of the t-mass values in the null distribution were marked as significant. For all significant clusters, we report the exact p-value (2 decimals) and the time range where the 95%-CIs of the observed data differ from zero (the longest deviation was reported). Note that we do not report the time range of the cluster itself, because cluster-level significance establishes only that an effect exists, not where it is situated in time (Sassenhagen & Draschkow, 2019). Since the procedure only tested for positive deviations from zero, time series with suspected negative deviations were sign-reversed.

## Results

### Behavioral Performance

All participants completed the task with an average accuracy (percentage of correctly identified relevant probes as mis-/matching the VWM Item) of *M* = 94.44% across all trials (*SD* = 4.90%). On average, participants performed similarly for both cues, with 94.89% (*SD* = 5.53%) for Cue 1 and 93.96% (*SD* = 4.95%) for Cue 2.

### RIFT reveals anticipatory early visual processing enhancements prior to relevant visual events

First, we addressed the main question by testing for pro-active neural responses before probe onset. To this end we ensured 60 Hz RIFT coherence was observed strongest in the posterior electrodes (Supplementary Figure 1). To determine whether visual processing was enhanced in anticipation of relevant compared to irrelevant visual events, we investigated the level of RIFT coherence before probe presentation.

In the interval before Probe 1 presentation, 60Hz RIFT coherence was higher when Probe 1 was cued to be relevant versus when it was cued to be irrelevant (permutation test, p = .04), supporting our hypothesis (Figure 2); 95%-CIs [2.00, 2.40]s. Similarly, RIFT coherence was higher before Probe 2 presentation when Probe 2 was cued as relevant compared to irrelevant (permutation test, p < .001), again supporting our hypothesis (Figure 2); 95%-CIs [3.57, 4.821]s. The average RIFT difference between relevant and irrelevant probes (i.e., Probe 1 & 2 relevant versus irrelevant) across significant intervals was localized in posterior electrodes. Interestingly, these differences in RIFT coherence did not persist beyond the anticipatory period. That is, the difference in RIFT responses between relevant and irrelevant conditions disappeared shortly after probe presentation.

**Figure 2:**
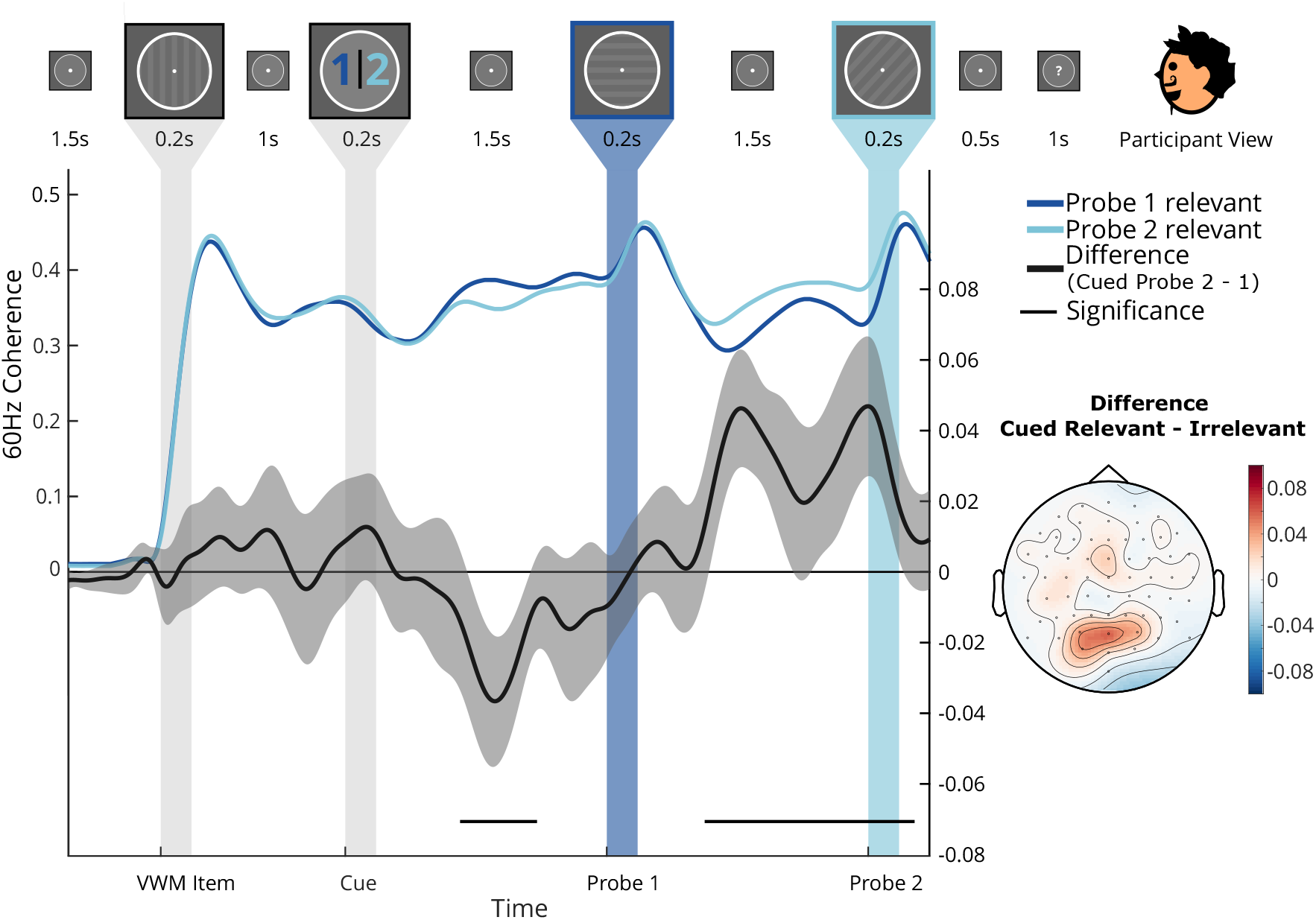
Anticipating a relevant (compared to irrelevant) probe enhanced visual processing *before* probe presentation. The top row depicts the time course of events as seen by participants. Average RIFT coherence (quantifying early visual processing; left y-axis) over time, separated for trials in which Probe 1 (dark blue) or Probe 2 (light blue) was relevant. The paired difference between conditions (right y-axis) is depicted in black. Shaded area: 95%-CI. Horizontal black line: significance according to a permutation-based cluster analysis, *p* <.05. Topography: The average RIFT coherence difference between relevant and irrelevant probes (i.e., Probe 1 & 2 relevant versus irrelevant) across significant intervals.

### Enhanced response to relevant probes

Next, we analyzed re-active neural responses to the probes to confirm that participants engaged with the temporal structure of the task. We expected that relevant probes would evoke a stronger univariate response than irrelevant probes. Indeed, we found an overall stronger EEG response to relevant visual events compared to irrelevant visual events, for both Probe 1 and Probe 2 (see Figure 3; t-test, p_adj_ < .05; permutation test_Probe1_, p < .001; 95%-CIs [0.17, 1.20]s; permutation test_Probe2_, p < .001; 95%-CIs [0.28 0.70]s).

**Figure 3:**
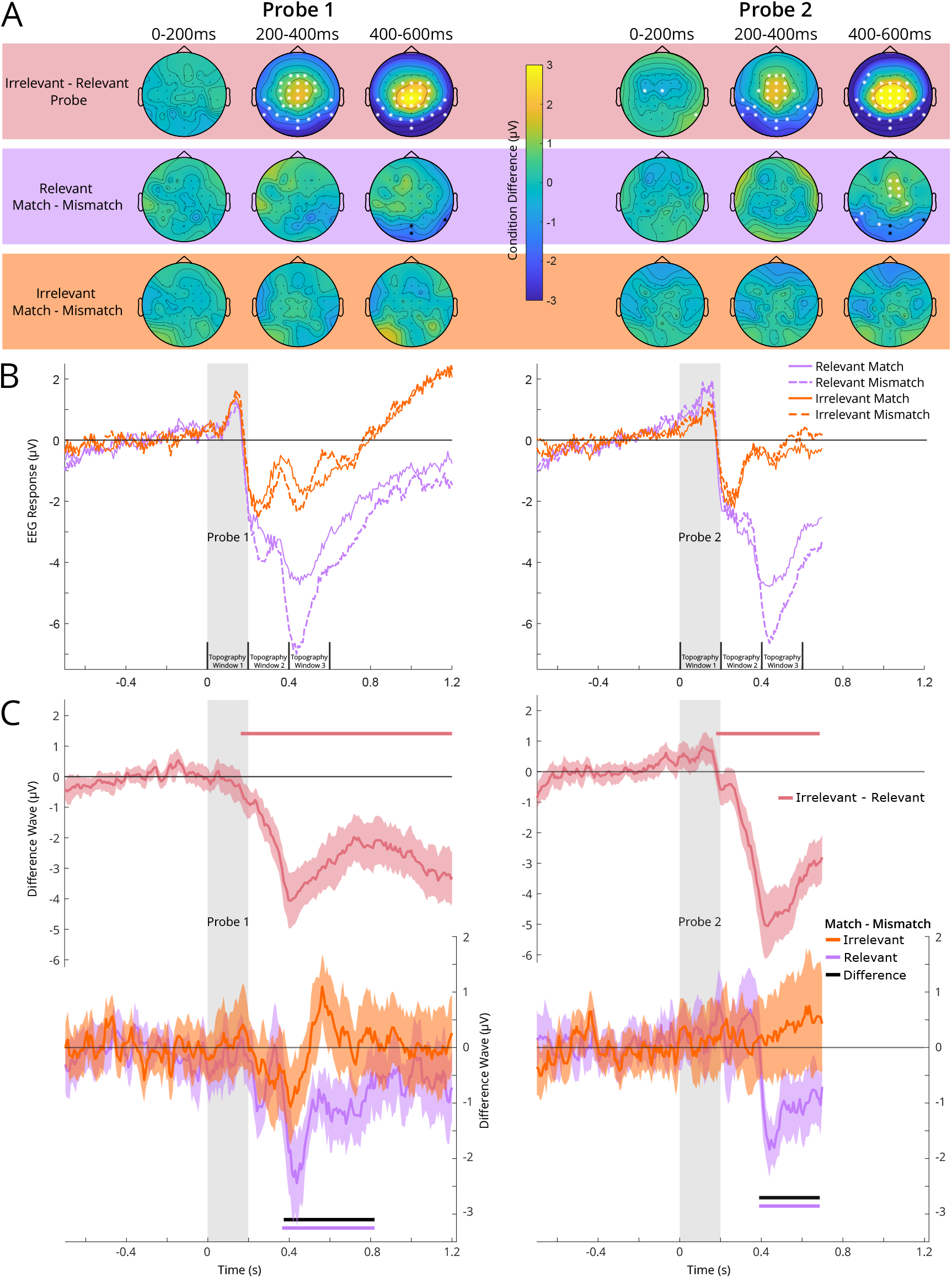
Univariate responses (i.e., event-related potentials) to Probe 1 and 2, for the four different conditions of interest (relevant and irrelevant, matching and mismatching probes). Panel A depicts whole-scalp topographies of the difference between relevant and irrelevant probes (top), and the difference between matching and mismatching probes when they are relevant (middle) and when they are irrelevant (bottom). Asterisks denote signficant channels according to a two-sided t-test (FDR corrected for multiple comparisons, *p* <.05). Panel B depicts the full time course of the event-related potential (ERP) for a selection of posterior electrodes (based on the whole-scalp topographies in Panel A; black asterisks), averaged across participants. Panel C depicts the difference between relevant and irrelevant time courses (top), and the difference between matching and mismatching time courses, separately for relevant and irrelevant probes (bottom). Shaded areas depict the 95%-CIs across participants. Horizontal thick lines denote significant ERP differences according to a permutation-based cluster analysis, *p* <.05.

Finally, effective use of temporal expectations during memory-guided visual selection of memory-matching and memory-mismatching visual events (i.e., blue highway exit signs versus white traffic signs) should evoke differential responses when they are relevant (i.e., in 5 km), but not when they are irrelevant (i.e., right now). Indeed, memory-matching and memory-mismatching events evoked a differential response when they were relevant (see Figure 3; t-test, p_adj_ < .05; permutation test_Probe1_, p < .001; 95%-CIs [.38 .80]s; permutation test_Probe2_, p < .001; 95%-CIs [.41, .70]s), but not when they were irrelevant (t-test, p_adj_ < .05), in posterior electrodes. Crucially, this differential response to relevant memory-matching versus memory-mismatch visual events was distinct from the differential (match/mismatch) response to irrelevant visual events (Figure 3C, Match - Mismatch: Difference; permutation test_Probe1_, p = .002; 95%-CIs [.38 .70]s; permutation test_Probe2_, p = .002; 95%-CIs [.41 .64]s)). This was the case for Probe 1 as well as for Probe 2. Thus to summarize: relevant visual events reliably evoked a stronger re-active response than irrelevant visual events. This response was only modulated by memory-content when the probe was cued as relevant.

## Discussion

We examined the dynamics of memory-guided visual selection in the face of relevant and irrelevant visual events. The results demonstrate that event relevance affects visual processing during memory-guided selection. Specifically, anticipating relevant — rather than irrelevant — visual events (1) increases RIFT coherence prior to the event onset, and subsequently (2a) increases univariate responses to the events themselves, and (2b) increases the response difference between memory-matching and memory-mismatching events. In sum, the anticipated (ir)relevance of a visual event (1) pro-actively enhances early visual processing, and (2) re-actively modulates the neural response to the visual event itself. Anticipating the (ir)relevance of upcoming events, then, enables the visual system to prepare for efficient memory-guided visual selection.

### Enhanced visual processing prior to relevant visual events

We hypothesized that the temporal structure of the task would allow observers to change the intensity of early visual processing over time, depending on when the relevant visual event was expected to occur. RIFT tracked anticipatory changes in visual processing (before event onset) with high temporal resolution. We found increased RIFT coherence preceding relevant compared to irrelevant probes, despite visual stimulation being constant (i.e., same temporal structure and balanced probe orientations). This was observed in the period preceding both probes, reflected in a mirrored difference wave (cued Probe 1 - cued Probe 2) across the duration of a trial. Specifically, when the first probe was relevant for the memory task, there was a relative increase in the RIFT response before the first probe, and a subsequent decrease before the second probe; conversely, when the second probe was relevant, there was a relative decrease in the RIFT response before the first probe, followed by an increase before the second probe. Our findings show that early visual processing can be flexibly up- and down-regulated over time, in anticipation of relevant visual events, relative to irrelevant visual events. These pro-active changes in early visual processing emerged well (approximately up to one second) before the actual stimulus was presented.

It is noteworthy that memory load differed in the period before Probe 1 and the period before Probe 2 due to the task design. Before Probe 1, participants had to maintain the memory item in both relevancy conditions, but in the delay period preceding Probe 2, participants could drop the memory item if Probe 1 was relevant (immediately after completing the comparison). The larger RIFT response preceding a relevant probe compared to an irrelevant probe in the second delay period (i.e., Cue 2 versus Cue 1) could therefore reflect a difference in memory load rather than a difference in relevance. This caveat similarly applies to the re-active responses to Probe 2. Critically, however, all key results were not only observed for Probe 2, but also for Probe 1, where memory load was identical between conditions (Figure 2; Figure 3). Therefore, both the pro-active RIFT response and the re-active univariate responses were uniquely modulated by relevance, and not confounded by differences in task demands.

Based on the present results alone, we cannot determine the absolute direction of the visual processing changes. Observers may up-regulate early visual processing in anticipation of relevant visual events (the cued item) or down-regulate early visual processing in anticipation of irrelevant visual events (the uncued item), or both. Future studies, including a condition without visual events (i.e., a baseline), may be needed to arbitrate between these possibilities definitively. It is worth noting that it is difficult to establish a true baseline of visual processing over time (even without stimulus presentations, participants will form expectations about upcoming visual input, e.g., Hazard rates). An alternative approach would be to reduce the formation (or usefulness) of expectations by making the onset time of the target unpredictable.

The present results do align with previous behavioral and neurophysiological research, which has identified a diverse range of processing benefits for inputs that can be anticipated (Bertelson & Boons, 1960; Coull & Nobre, 1998; Heuer & Rolfs, 2022; Klemmer, 1956; Miniussi et al., 1999; Nobre & Van Ede, 2018; Rohenkohl et al., 2012; Vangkilde et al., 2012). In most of these studies, there was no "distracting" (or irrelevant) visual event that required suppression. Typically, participants are cued to expect either an imminent relevant or a distant relevant target. It is plausible that participants up-regulate processing in anticipation of imminent relevant visual events rather than (only) down-regulate processing in anticipation of distant relevant visual events. This suggests that anticipatory effects observed in the present study — at least partly — reflect up-regulation of visual processing in the face of relevant visual events (and cannot be fully explained by the suppression of irrelevant visual events). Furthermore, the results extend previous findings investigating the anticipated (ir)relevance of visual events using visible frequency tagging (SSVEPs), which already hinted at an increase in visual processing in anticipation of relevant compared to irrelevant probes (Denison et al., 2024), but could not definitively distinguish between pro-active and re-active effects due to the visibility of the tagging (while the frequency tag is irrelevant to the task, the visible changes on screen still act as an exogenous cue on visual attention and possibly interfering with top-down control).

At face value, the observed pro-active visual processing changes are similar to earlier work on temporal expectations showing anticipatory modulations in visual processing in the absence of a memory task (Samaha et al., 2015; Van Ede et al., 2020). However, it remains an open question whether and how anticipatory changes in visual processing depend on task context; particularly whether or not visual information is concurrently maintained in working memory. Future studies should thus directly compare visual processing preparation during memory-guided visual selection to visual processing during pure perceptual tasks.

Crucially, the present study expands on this previous literature on temporal expectations by measuring how anticipating the (ir)relevance of a visual event affects early visual processing in particular. RIFT is interpreted to reflect neural excitability of the early visual cortex (Duecker et al., 2021, 2025), after Studies using RIFT with MEG have localized the source of the RIFT signal to primary and secondary visual cortex (Duecker et al., 2021, 2025; Schneider et al., 2023; Zhigalov & Jensen, 2020). In addition, because RIFT is experimenter-induced (and because the measurement of the RIFT response relies on the known phase of the oscillatory response), the RIFT response can be isolated from other, non-phase-aligned oscillations in the neural signal, even when occurring in similar frequency ranges (e.g., eye-movement-induced gamma-band responses (Yuval-Greenberg et al., 2008)). Previous research using multiple RIFT frequencies (i.e., 60Hz and 64Hz) has shown that when phase aligning (phase-shuffled) trials for one frequency (e.g., 60Hz), there is no coherence response for the other (e.g., 64Hz; Arora, Husta, et al., 2025; Wang, Arora, et al., 2025). In contrast, the exact relationship of endogenous responses (e.g., alpha oscillations) to visual processing is debated (Brickwedde et al., 2025; Zhigalov & Jensen, 2020). Therefore, previous evidence relying on endogenous responses to conclude about anticipatory visual processing changes is equally ambiguous. Here we show, for the first time, that temporal expectations modulate specifically early visual processing in the period preceding (ir)relevant stimuli. These anticipatory changes in early visual processing may lie at the basis of behavioral and neural biases observed in extant studies manipulating temporal expectations.

### Enhanced neural response to relevant visual events

Moving beyond the anticipatory period, we also asked how relevance influences the neural (re-active) responses to the visual events themselves. We found that relevant visual events evoke a stronger neural response than irrelevant visual events. Importantly, relevant and irrelevant visual events did not differ systematically in their visual features and, therefore, do not underlie the observed response differences. Overall then, these results align with existing work observing stronger (re-active) neural responses to targets presented at (cued) relevant compared to irrelevant spatial locations (Hillyard et al., 1998; Mangun & Hillyard, 1990), and time points (Correa et al., 2006; Miniussi et al., 1999).

Within the context of VWM-guided selection, the preparatory enhancement of visual processing may facilitate perceptual comparisons between the relevant visual events (what is seen) and VWM content (what is searched for). Thus, we hypothesized that the response difference between memory-matching and memory-mismatching visual events (i.e., the object one is looking for versus other objects) is larger when the item is currently relevant, compared to when it is not. In line with this hypothesis, we found dissociable univariate responses to (i.e., ERP differences between) memory-matching and memory-mismatching visual events when probes were relevant for the memory task, but not when they were irrelevant. Thus, temporal expectations are used to regulate the memory-based differentiation between target and distractor objects.

This finding makes intuitive sense because — both in our experiment and in daily life — target and distractor objects require different behavioral actions (exit the highway or not) when they are relevant, but not when they are irrelevant. The present findings add to a growing literature investigating how memory-based modulations of visual processing depend on the attentional state of the visual working memory content. Previous literature has shown that currently relevant memory items bias concurrent visual processing more strongly than currently irrelevant memory items (Olivers et al., 2011; Wang, Chota, et al., 2025). This line of research, however, mainly compared how multiple parallel memory items in different attentional (or: relevancy) states influence perception. In this case, however, we explicitly studied how changes in the relevance of a single memory item influenced concurrent visual processing. This may be more similar to real-world situations, in which relevancy changes over time, and observers typically do not load more than one item in memory (Draschkow et al., 2021; Qing et al., 2025; Sahakian et al., 2023; Van der Stigchel et al., 2019). Here we show that as the relevance of a single memory item for behavior waxes and wanes, its influence on visual selection follows.

An exploratory univariate analysis of the whole trial (see Supplementary Figure 2) revealed baseline differences between relevance conditions already before probe onset; anticipating probes as relevant (compared to irrelevant) increased EEG voltage (significant only for Probe 2), thus mirroring the effects observed with RIFT. Tentatively, the (non-significant) pro-active change in EEG voltage before relevant (compared to irrelevant) probes could be regarded as a preparatory mechanism to enable a subsequent stronger re-active response to relevant probes.

### Anticipatory changes in memory representations

An outstanding question pertains to whether and how memory representations change in anticipation of an upcoming visual event. Indeed, in parallel to the observed enhancement of visual processing prior to relevant visual events, one may also expect an anticipatory sharpening of the visual working memory content. In support of this latter possibility, recent studies investigating the impact of temporal expectations on memory recall revealed that VWM content is more resistant to predictable than to unpredictable visual interference (Gresch et al., 2021; Van Ede et al., 2018). However, these studies did not directly measure the representational format or strength of the memory content in anticipation of the (relevant and irrelevant) visual events. More direct evidence comes from fMRI decoding studies, in which a cue indicated which of two items held in VWM would be relevant for the next of two consecutive recall tasks. These studies have shown that observers can indeed flexibly enhance relevant VWM representations in anticipation of a memory task (Christophel et al., 2018; LaRocque et al., 2017; Lewis-Peacock et al., 2012; Rose et al., 2016; Sahan et al., 2020; van Loon et al., 2018; Yu et al., 2020). In the context of visual search, recent fMRI decoding studies have shown that working memory representations of a search target — generated during search preparation — depend on contextual expectations (e.g., about the location of a target object, or the appearance of distractor objects; Gayet & Peelen, 2022; Lerebourg et al., 2024). These findings demonstrate that VWM contents can be strategically altered to match situational demands in a search task context. Relatedly, VWM representations can be dynamically adjusted (from a sensory to an action-oriented format) to meet current task demands (Henderson et al., 2022).

Together, these findings highlight the adaptability of working memory contents to the changing demands of dynamic environments. By extension, VWM representations likely also vary in response to the type of temporal expectations investigated. Unfortunately, EEG decoding methods thus far have been relatively unsuccessful in decoding the content of visual working memory over a prolonged period. Therefore, future work is needed (e.g., using combined fMRI-EEG) to directly test whether the changes in early visual processing established here are accompanied by a corresponding sharpening of the VWM content.

## Conclusion

In sum, when driving on a highway, you do not continuously and indiscriminately compare the visual input of the right side of the road with the highway exit sign that you are looking for. Instead, you prioritize visual input during the period in time when you expect the sign to appear. Specifically, when the anticipated highway exit is imminent, you preemptively enhance early visual processing, which may benefit the comparison between the visual information collected by your sensory organs and the visual representation of the exit sign held in memory. Taken together, anticipating the relevance of visual events enables the distribution of scarce attentional resources over time, in support of efficient memory-guided visual selection.

## Ethics approval

The study was conducted in accordance with the protocol approved by the Ethics Committee of the Faculty of Social and Behavioural Sciences at Utrecht University with reference number 863732 – HOMEOSTASIS (approved on 18/03/2020).

## Resource Availbility

### Lead Contact

Lasse Dietz Helmholtz Institute, Utrecht University 3584CS Utrecht, The Netherlands

### Materials Availability

Not applicable.

### Data and Code Availability

Preprocessed EEG, raw behavioral data, and the corresponding analysis are available as of the date of publication at the Open Science Framework: https://osf.io/7ve8m/.

### Author Contributions

: Conceptualization, L.D., C.S., S.V.d.S., S.C., and S.G.; data curation, L.D.; formal analysis, L.D., C.S., K.A., S.C., and S.G.; funding acquisition, S.C. and S.G.; methodology, L.D., C.S., K.A., S.C., and S.G.; project administration, L.D., C.S., S.V.d.S., S.C., and S.G.; software, L.D., K.A., and S.C.; supervision, C.S., S.V.d.S., S.C., and S.G.; visualization, L.D.; writing — original draft preparation, L.D. and S.G.; writing — review and editing, C.S., S.V.d.S., S.C., and S.G.

### Declaration of generative AI and AI-assisted technologies in the writing process

During the preparation of this work, the author(s) used ChatGPT in order to generate possible alternative sentence structures. After using this tool or service, the author(s) reviewed and edited the content as needed and take(s) full responsibility for the content of the publication.

## Supplemental Information

Document S1. Figures SupFig1-2.

## References

Arora, K., Gayet, S., Kenemans, J. L., Van Der Stigchel, S., & Chota, S. (2025). Dissociating external and internal attentional selection. iScience, 28(4), 112282. 10.1016/j.isci.2025.112282

Arora, K., Gayet, S., Kenemans, J. L., Van der Stigchel, S., & Chota, S. (2024). Rapid Invisible Frequency Tagging (RIFT) in a novel setup with EEG. bioRxiv, 2024–02.

Arora, K., Husta, C., Bouwkamp, F., Seijdel, N., Bai, S., Han, Q., Kenemans, L., Van der Stigchel, S., Gayet, S., Spaak, E., Chota, S., & Drijvers, L. (2025, October). A Collaborative Guide to Rapid Invisible Frequency Tagging (RIFT): Methods, Insights, and Recommendations. 10.31234/osf.io/edshx_v1

Benjamini, Y., & Hochberg, Y. (1995). Controlling the False Discovery Rate: A Practical and Powerful Approach to Multiple Testing. Journal of the Royal Statistical Society Series B: Statistical Methodology, 57(1), 289–300. 10.1111/j.2517-6161.1995.tb02031.x

Benjamini, Y., & Yekutieli, D. (2001). The control of the false discovery rate in multiple testing under dependency. Annals of statistics, 1165–1188.

Benjamini, Y., & Yekutieli, D. (2005). False Discovery Rate–Adjusted Multiple Confidence Intervals for Selected Parameters. Journal of the American Statistical Association, 100(469), 71–81. 10.1198/016214504000001907

Bertelson, P., & Boons, J.-P. (1960). Time Uncertainty and Choice Reaction Time. Nature, 187(4736), 531–532. 10.1038/187531a0

Brainard, D. H., & Vision, S. (1997). The psychophysics toolbox. Spatial vision, 10(4), 433–436.

Brickwedde, M., Limachya, R., Markiewicz, R., Sutton, E., Postzich, C., Shapiro, K., Jensen, O., & Mazaheri, A. (2025). Cross-modal interaction of Alpha Activity does not reflect inhibition of early sensory processing: A frequency tagging study using EEG and MEG. eLife, 14. 10.7554/eLife.106050.3

Çelik, Ü. G., Arora, K., Kenemans, J. L., Stigchel, S. V. d., Gayet, S., & Chota, S. (2025, October). Tracking attention using RIFT with a consumer-monitor setup. 10.1101/2025.10.03.680199

Christophel, T. B., Iamshchinina, P., Yan, C., Allefeld, C., & Haynes, J.-D. (2018). Cortical specialization for attended versus unattended working memory. Nature neuroscience, 21(4), 494–496.

Correa, Á., Lupiáñez, J., Madrid, E., & Tudela, P. (2006). Temporal attention enhances early visual processing: A review and new evidence from event-related potentials. Brain Research, 1076(1), 116–128. 10.1016/j.brainres.2005.11.074

Coull, J. T., & Nobre, A. C. (1998). Where and When to Pay Attention: The Neural Systems for Directing Attention to Spatial Locations and to Time Intervals as Revealed by Both PET and fMRI. The Journal of Neuroscience, 18(18), 7426– 7435. 10.1523/JNEUROSCI.18-18-07426.1998

Cravo, A. M., Rohenkohl, G., Santos, K. M., & Nobre, A. C. (2017). Temporal anticipation based on memory. Journal of cognitive neuroscience, 29(12), 2081–2089.

Cravo, A. M., Rohenkohl, G., Wyart, V., & Nobre, A. C. (2013). Temporal Expectation Enhances Contrast Sensitivity by Phase Entrainment of Low-Frequency Oscillations in Visual Cortex. The Journal of Neuroscience, 33(9), 4002–4010. 10.1523/JNEUROSCI.4675-12.2013

Denison, R. N., Tian, K. J., Heeger, D. J., & Carrasco, M. (2024). Anticipatory and evoked visual cortical dynamics of voluntary temporal attention. Nature Communications, 15(1), 9061. 10.1038/s41467-024-53406-y

Desimone, R., & Duncan, J. (1995). Neural mechanisms of selective visual attention.Annual review of neuroscience, 18(1), 193–222.

Downing, P. E. (2000). Interactions Between Visual Working Memory and Selective Attention. Psychological Science (0956-7976), 11(6). 10.1111/1467-9280.00290

Draschkow, D., Kallmayer, M., & Nobre, A. C. (2021). When natural behavior engages working memory. Current Biology, 31(4), 869–874.

Duecker, K., Gutteling, T. P., Herrmann, C. S., & Jensen, O. (2021). No evidence for entrainment: Endogenous gamma oscillations and rhythmic flicker responses coexist in visual cortex. Journal of Neuroscience, 41(31), 6684–6698.

Duecker, K., Shapiro, K. L., Hanslmayr, S., Griffiths, B. J., Pan, Y., Wolfe, J., & Jensen, O. (2024, October). Guided Visual Search is associated with a feature-based priority map in early visual cortex. 10.1101/2024.10.03.616568

Duecker, K., Shapiro, K. L., Hanslmayr, S., Griffiths, B. J., Pan, Y., Wolfe, J. M., & Jensen, O. (2025). Guided visual search is associated with target boosting and distractor suppression in early visual cortex. Communications Biology, 8(1), 912.

Duncan, J., & Humphreys, G. W. (1989). Visual search and stimulus similarity. Psychological review, 96(3), 433.

Eimer, M. (2014). The neural basis of attentional control in visual search. Trends in cognitive sciences, 18(10), 526–535.

Ferrante, O., Zhigalov, A., Hickey, C., & Jensen, O. (2023). Statistical Learning of Distractor Suppression Downregulates Prestimulus Neural Excitability in Early Visual Cortex. The Journal of Neuroscience, 43(12), 2190–2198. 10.1523/JNEUROSCI.1703-22.2022

Gayet, S., Guggenmos, M., Christophel, T. B., Haynes, J.-D., Paffen, C. L., Van Der Stigchel, S., & Sterzer, P. (2017). Visual Working Memory Enhances the Neural Response to Matching Visual Input. The Journal of Neuroscience, 37(28), 6638–6647. 10.1523/JNEUROSCI.3418-16.2017

Gayet, S., Paffen, C. L. E., & Van Der Stigchel, S. (2013). Information Matching the Content of Visual Working Memory Is Prioritized for Conscious Access. Psychological Science, 24(12), 2472–2480. 10.1177/0956797613495882

Gayet, S., & Peelen, M. V. (2022). Preparatory attention incorporates contextual expectations. Current Biology, 32(3), 687–692.

Gresch, D., Boettcher, S. E., Van Ede, F., & Nobre, A. C. (2021). Shielding working-memory representations from temporally predictable external interference. Cognition, 217, 104915.

Harrison, S. A., & Tong, F. (2009). Decoding reveals the contents of visual working memory in early visual areas. Nature, 458(7238), 632–635.

Henderson, M. M., Rademaker, R. L., & Serences, J. T. (2022). Flexible utilization of spatial-and motor-based codes for the storage of visuo-spatial information. Elife, 11, e75688.

Heuer, A., & Rolfs, M. (2022). A direct comparison of attentional orienting to spatial and temporal positions in visual working memory. Psychonomic Bulletin & Review, 29(1), 182–190. 10.3758/s13423-021-01972-3

Hillyard, S. A., Vogel, E. K., & Luck, S. J. (1998). Sensory gain control (amplification) as a mechanism of selective attention: Electrophysiological and neuroimaging evidence (G. W. Humphreys, J. Duncan, & A. Treisman, Eds.). Philosophical Transactions of the Royal Society of London. Series B: Biological Sciences, 353(1373), 1257–1270. 10.1098/rstb.1998.0281

Hustá, C., Meyer, A., & Drijvers, L. (2025). Using rapid invisible frequency tagging (RIFT) to probe the neural interaction between representations of speech planning and comprehension. Neurobiology of Language, 1–16.

Jin, W., Nobre, A. C., & van Ede, F. (2020). Temporal expectations prepare visual working memory for behavior. Journal of Cognitive Neuroscience, 32(12), 2320–2332.

Kastner, S., & Ungerleider, L. G. (2001). The neural basis of biased competition in human visual cortex. Neuropsychologia, 39(12), 1263–1276. 10.1016/S0028-3932(01)00116-6

Klemmer, E. T. (1956). Time uncertainty in simple reaction time. Journal of Experimental Psychology, 51(3), 179–184.

LaRocque, J. J., Riggall, A. C., Emrich, S. M., & Postle, B. R. (2017). Within-category decoding of information in different attentional states in short-term memory. Cerebral Cortex, 27(10), 4881–4890.

Lerebourg, M., De Lange, F. P., & Peelen, M. V. (2024). Attentional guidance through object associations in visual cortex. Science Advances, 10(41). 10.1126/sciadv.ado6226

Lewis-Peacock, J. A., Drysdale, A. T., Oberauer, K., & Postle, B. R. (2012). Neural Evidence for a Distinction between Short-term Memory and the Focus of Attention. Journal of Cognitive Neuroscience, 24(1), 61–79. 10.1162/jocn_a_00140

Mangun, G., & Hillyard, S. (1990). Allocation of visual attention to spatial locations: Tradeoff functions for event-related brain potentials and detection performance. Perception & Psychophysics, 47(6), 532–550. 10.3758/BF03203106

Maris, E., & Oostenveld, R. (2007). Nonparametric statistical testing of EEG- and MEG-data. Journal of Neuroscience Methods, 164(1), 177–190. 10.1016/j.jneumeth.2007.03.024

Miniussi, C., Wilding, E. L., Coull, J. T., & Nobre, A. C. (1999). Orienting attention in time: Modulation of brain potentials. Brain, 122(8), 1507–1518.

Nobre, A. C., & Van Ede, F. (2018). Anticipated moments: Temporal structure in attention. Nature Reviews Neuroscience, 19(1), 34–48.

Olivers, C. N., Meijer, F., & Theeuwes, J. (2006). Feature-based memory-driven attentional capture: Visual working memory content affects visual attention. Journal of Experimental Psychology: Human Perception and Performance, 32(5), 1243.

Olivers, C. N., Peters, J., Houtkamp, R., & Roelfsema, P. R. (2011). Different states in visual working memory: When it guides attention and when it does not. Trends in cognitive sciences, 15(7), 327–334.

Olmos-Solis, K., van Loon, A. M., Los, S. A., & Olivers, C. N. (2017). Oculomotor measures reveal the temporal dynamics of preparing for search. Progress in brain research, 236, 1–23.

Oostenveld, R., Fries, P., Maris, E., & Schoffelen, J.-M. (2011). FieldTrip: Open Source Software for Advanced Analysis of MEG, EEG, and Invasive Electrophysiological Data. Computational Intelligence and Neuroscience, 2011, 1–9. 10.1155/2011/156869

Pan, Y., Frisson, S., Federmeier, K. D., & Jensen, O. (2022, September). Parafoveal semantic integration in natural reading. 10.1101/2022.09.26.509511

Pelli, D. G. (1997). The VideoToolbox software for visual psychophysics: Transforming numbers into movies. Spatial vision, 10(4), 437–442.

Qing, T., Strauch, C., Van Maanen, L., & Van Der Stigchel, S. (2025). Shifting reliance between the internal and external world: A meta-analysis on visual-working memory use. Psychonomic Bulletin & Review, 32(3), 1118–1130. 10.3758/s13423-024-02623-z

Rohenkohl, G., Cravo, A. M., Wyart, V., & Nobre, A. C. (2012). Temporal expectation improves the quality of sensory information. Journal of Neuroscience, 32(24), 8424–8428.

Rose, N. S., LaRocque, J. J., Riggall, A. C., Gosseries, O., Starrett, M. J., Meyering, E. E., & Postle, B. R. (2016). Reactivation of latent working memories with transcranial magnetic stimulation. Science, 354(6316), 1136–1139. 10.1126/science.aah7011

Sahakian, A., Gayet, S., Paffen, C. L., & Van Der Stigchel, S. (2023). Mountains of memory in a sea of uncertainty: Sampling the external world despite useful information in visual working memory. Cognition, 234, 105381. 10.1016/j.cognition.2023.105381

Sahan, M. I., Sheldon, A. D., & Postle, B. R. (2020). The neural consequences of attentional prioritization of internal representations in visual working memory. Journal of Cognitive Neuroscience, 32(5), 917–944.

Samaha, J., Bauer, P., Cimaroli, S., & Postle, B. R. (2015). Top-down control of the phase of alpha-band oscillations as a mechanism for temporal prediction. Proceedings of the National Academy of Sciences, 112(27), 8439–8444. 10.1073/pnas.1503686112

Sassenhagen, J., & Draschkow, D. (2019). Cluster-based permutation tests of MEG/EEG data do not establish significance of effect latency or location. Psychophysiology, 56(6). 10.1111/psyp.13335

Schneider, M., Tzanou, A., Uran, C., & Vinck, M. (2023). Cell-type-specific propagation of visual flicker. Cell Reports, 42(5), 112492. 10.1016/j.celrep.2023.112492

Serences, J. T., Ester, E. F., Vogel, E. K., & Awh, E. (2009). Stimulus-Specific Delay Activity in Human Primary Visual Cortex. Psychological Science, 20(2), 207–214. 10.1111/j.1467-9280.2009.02276.x

Silvis, J. D., & Van Der Stigchel, S. (2014). How memory mechanisms are a key component in the guidance of our eye movements: Evidence from the global effect. Psychonomic Bulletin & Review, 21(2), 357–362. 10.3758/s13423-013-0498-9

Soto, D., Hodsoll, J., Rotshtein, P., & Humphreys, G. W. (2008). Automatic guidance of attention from working memory. Trends in cognitive sciences, 12(9), 342–348.

Van der Stigchel, S., Schut, M., & Somai, R. (2019). Evidence for the world as an external memory: A trade-off between internal and external visual memory storage. Journal of Vision, 19(10), 78–78.

Van Ede, F., Chekroud, S. R., Stokes, M. G., & Nobre, A. C. (2018). Decoding the influence of anticipatory states on visual perception in the presence of temporal distractors. Nature communications, 9(1), 1449.

Van Ede, F., Niklaus, M., & Nobre, A. C. (2017). Temporal Expectations Guide Dynamic Prioritization in Visual Working Memory through Attenuated Oscillations. The Journal of Neuroscience, 37(2), 437–445. 10.1523/JNEUROSCI.2272-16.2016

Van Ede, F., Rohenkohl, G., Gould, I., & Nobre, A. C. (2020). Purpose-Dependent Consequences of Temporal Expectations Serving Perception and Action. The Journal of Neuroscience, 40(41), 7877–7886. 10.1523/JNEUROSCI.1134-20.2020

Vangkilde, S., Coull, J. T., & Bundesen, C. (2012). Great expectations: Temporal expectation modulates perceptual processing speed. Journal of Experimental Psychology: Human Perception and Performance, 38(5), 1183.

van Loon, A. M., Olmos-Solis, K., Fahrenfort, J. J., & Olivers, C. N. (2018). Current and future goals are represented in opposite patterns in object-selective cortex. elife, 7, e38677.

Walter, W. G., Cooper, R., Aldridge, V. J., McCallum, W. C., & Winter, A. L. (1964). Contingent negative variation: An electric sign of sensori-motor association and expectancy in the human brain. nature, 203(4943), 380–384.

Wang, D., Arora, K., Theeuwes, J., Stigchel, S. V. d., Gayet, S., & Chota, S. (2025, August). Dynamic competition between bottom-up saliency and top-down goals in early visual cortex. 10.1101/2025.08.22.671530

Wang, D., Chota, S., Xu, L., der Stigchel, S. V., & Gayet, S. (2025). The priority state of items in visual working memory determines their influence on early visual processing. Consciousness and Cognition, 127, 103800. 10.1016/j.concog.2024.103800

Wilsch, A., Henry, M. J., Herrmann, B., Herrmann, C. S., & Obleser, J. (2018). Temporal Expectation Modulates the Cortical Dynamics of Short-Term Memory. The Journal of Neuroscience, 38(34), 7428–7439. 10.1523/JNEUROSCI.2928-17.2018

Wolfe, J. M. (1994). Guided Search 2.0 A revised model of visual search. Psychonomic Bulletin & Review, 1(2), 202–238. 10.3758/bf03200774

Wolfe, J. M., & Horowitz, T. S. (2004). What attributes guide the deployment of visual attention and how do they do it? Nature reviews neuroscience, 5(6), 495–501.

Yu, Q., Teng, C., & Postle, B. R. (2020). Different states of priority recruit different neural representations in visual working memory. PLoS biology, 18(6), e3000769.

Yuval-Greenberg, S., Tomer, O., Keren, A. S., Nelken, I., & Deouell, L. Y. (2008). Transient induced gamma-band response in EEG as a manifestation of miniature saccades. Neuron, 58(3), 429–441.

Zhigalov, A., Herring, J. D., Herpers, J., Bergmann, T. O., & Jensen, O. (2019). Probing cortical excitability using rapid frequency tagging. NeuroImage, 195, 59–66. 10.1016/j.neuroimage.2019.03.056

Zhigalov, A., & Jensen, O. (2020). Alpha oscillations do not implement gain control in early visual cortex but rather gating in parieto-occipital regions. Human Brain Mapping, 41(18), 5176–5186. 10.1002/hbm.25183

Zokaei, N., Board, A. G., Manohar, S. G., & Nobre, A. C. (2019). Modulation of the pupillary response by the content of visual working memory. Proceedings of the National Academy of Sciences, 116(45), 22802–22810. 10.1073/pnas.1909959116

