## Supplemental File 1 for "Anticipated Relevance Prepares Visual Processing for Efficient Memory-Guided Selection"

* Shared authorship

### RIFT Topographies Per Participant

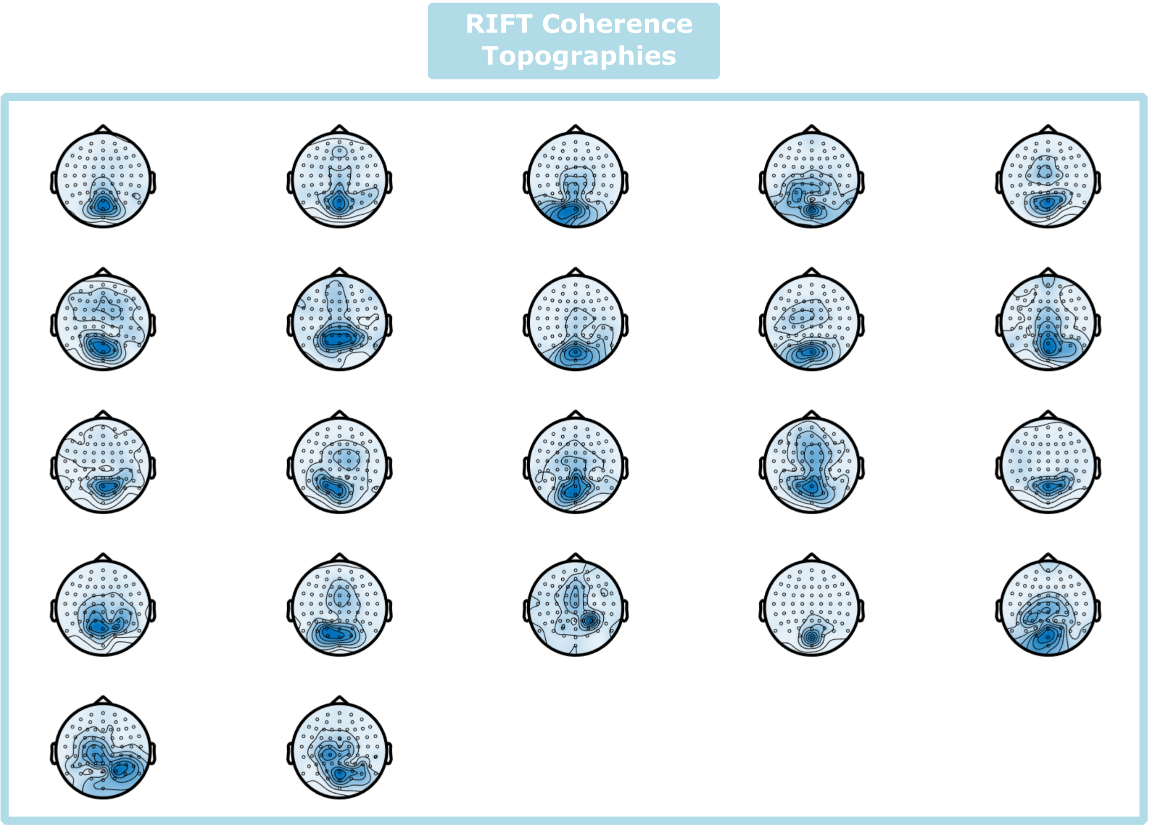


Supplementary Figure 1: Topographic overview of 60 Hz RIFT coherence strength per participant.

### Increased EEG-voltage when anticipating relevant probes

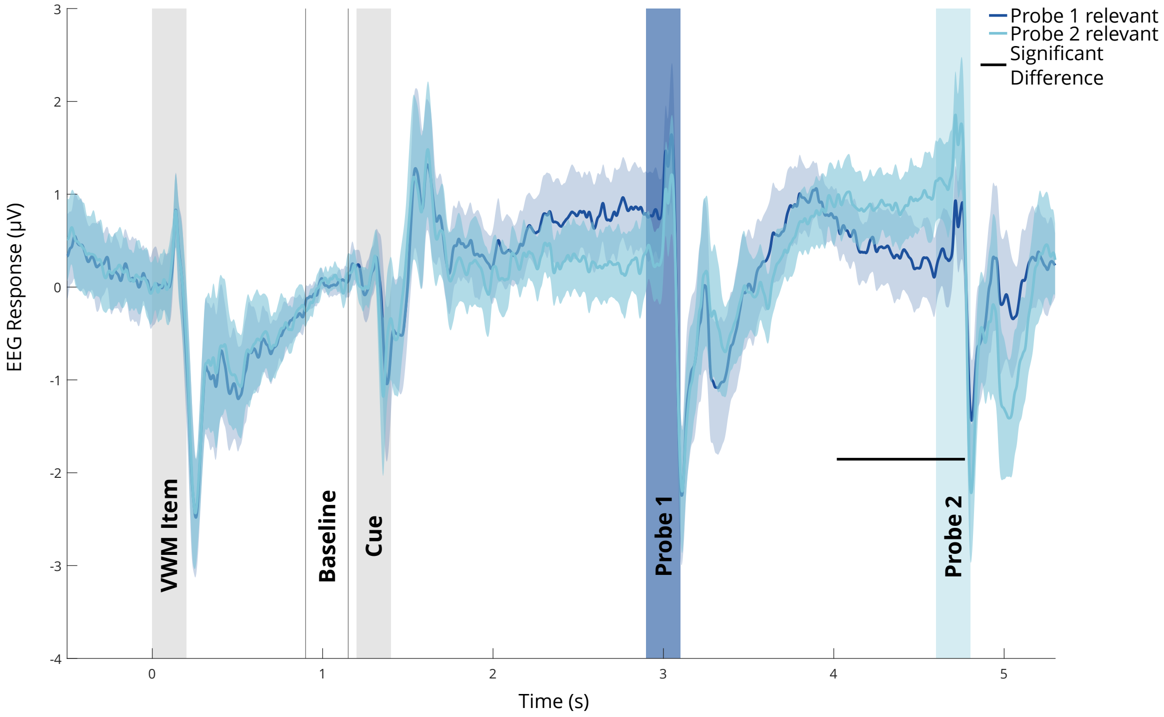


Supplementary Figure 2: Anticipating probes as relevant – compared to irrelevant – increased EEG voltage (significant only for Probe 2) . Depicted: Average (across participants) EEG response over time, separated for trials in which Probe 1 (dark blue) or Probe 2 (light blue) was relevant. Baseline correction (two vertical black lines) was applied before the cue (900-1150ms). Shaded area: 95%-CI. Horizontal black line: significance according to a permutation-based cluster analysis, $p<.05$.
